# Assessing performance of pathogenicity predictors using clinically-relevant variant datasets

**DOI:** 10.1101/2020.02.06.937169

**Authors:** Adam C Gunning, Verity Fryer, James Fasham, Andrew H Crosby, Sian Ellard, Emma Baple, Caroline F Wright

## Abstract

**Purpose:** Pathogenicity predictors are an integral part of genomic variant interpretation but, despite their widespread usage, an independent validation of performance using a clinically-relevant dataset has not been undertaken.

**Methods:** We derive two validation datasets: an “open” dataset containing variants extracted from publicly-available databases, similar to those commonly applied in previous benchmarking exercises, and a “clinically-representative” dataset containing variants identified through research/diagnostic exome and diagnostic panel sequencing. Using these datasets, we evaluate the performance of three recently developed meta-predictors, REVEL, GAVIN and ClinPred, and compare their performance against two commonly used *in silico* tools, SIFT and PolyPhen-2.

**Results:** Although the newer meta-predictors outperform the older tools, the performance of all pathogenicity predictors is substantially lower in the clinically-representative dataset. Using our clinically-relevant dataset, REVEL performed best with an area under the ROC of 0.81. Using a concordance-based approach based on a consensus of multiple tools reduces the performance due to both discordance between tools and false concordance where tools make common misclassification. Analysis of tool feature usage may give an insight into the tool performance and misclassification.

**Conclusion:** Our results support the adoption of meta-predictors over traditional *in silico* tools, but do not support a consensus-based approach as recommended by current variant classification guidelines.

## 1. INTRODUCTION

As the scale of genomic sequencing continues to increase, the classification of rare genomic variants is becoming the primary bottle-neck in the diagnosis of rare monogenic disorder. Guidelines published by the American College of Medical Genetics (ACMG) in 2016^1^ have helped bring consistency to variant classification and have been followed by a number of regional and disorder-specific publications^2–4^. Common to all guidelines is the recommendation of the use of *in silico* prediction tools to aid in the classification of missense variants. *In silico* prediction tools are algorithms designed to predict the functional impact of variation, usually missense changes caused by single nucleotide variants (SNVs). Though originally designed for the prioritisation of research variants^5^, the tools are used routinely in clinical diagnostics during variant classification. The tools integrate a number of features in order assess the impact of a variant on protein function^6^. Initially, inter-species conservation formed the bulk of the predictions, with some additional functional information, such as substitution matrices of physicochemical distances of amino acids (such as Grantham^7^ or PAM^8^), and data derived from a limited number of available X-ray crystallographic structures^9^. Since the development of the first *in silico* prediction tools over a decade ago^5,9^, large-scale experiments such as the ENCODE project^10^ have generated huge amounts of functional data, and we now also have access to large-scale databases of clinical and neutral variation^11–13^. These additional sources of data have led to an explosion of new *in silico* prediction algorithms^14–16^ that purport to increase accuracy.

However, the large increase in the number of predictors integrated into classification algorithms has raised concerns about overfitting^17,18^. Overfitting occurs when the prediction algorithm is trained on superfluous data or features that are irrelevant to the prediction outcome^18^. While it may appear that an increasingly large feature list leads to improvements in prediction, random variability within the training dataset may actually result in decreased accuracy when applied to a novel dataset. Overfitting can be mitigated through the use of increasingly large training datasets, and the usage of online variant databases, such as the genome aggregation database (gnomAD)^19^ and ClinVar^12^, allows for sufficiently large training datasets. Additionally, reliance on additional information – such as protein functional data and allele frequency data such as from gnomAD^19^ – may be contrary to the standard assumptions of variant classification methodology, namely that each dataset is independent and applied only once during classification.

Current ACMG guidelines recommend the use of a concordance-based approach, where a number of prediction algorithms are used, and evidence is applied only when there is agreement between tools. There is no guidance on which *in silico* tools should be used, how many, or on what constitutes a consensus, and this ambiguity allows for inconsistencies in the application of this piece of evidence across clinical laboratories. Studies have previously identified the limitations of applying a strict binary consensus-based approach^20^. In response, multiple groups^14–16^ have created meta-predictors; tools which integrate information from a large number of sources into a machine-learning algorithm. These tools thereby adhere to the principle of the consensus-based model suggested by ACMG without the onerous task of determining tool concordance, and reduce discordance when increasingly large numbers of tools are utilised. Unlike a manual consensus-based model, where tools are weighted equally, meta-predictors are able to apply weighting to features in order to maximise accuracy.

In order to evaluate the accuracy of *in silico* prediction tools, precompiled variant datasets such as VariBench^21^ have been designed to aid in training and benchmarking of pathogenicity predictors. However, the use of standardised datasets may introduce inherent biases into prediction algorithms, resulting in false concordance. Typically, prediction software is trained using machine-learning algorithms, and assessed using variants available from large online public databases ^5,6,9,10,14–16,22^ such as ExAC/gnomAD, ClinVar^12^, and SwissProt^23^. It has been previously shown that prediction algorithms have variable performance when applied to different datasets^6,22,24,25^, and therefore the use of variant datasets derived from online public databases may not be representative of the performance of tools when applied in a clinical setting. While studies emphasise the use of ‘neutral’ variation, the output from a modern next-generation sequencing pipeline is generally far from neutral, and includes a large number of variant filtering steps in order to reduce the burden of manual variant assessment^26^.

Here we evaluate and compare the performance of two traditional *in silico* pathogenicity prediction tools commonly used for clinical variant interpretation (SIFT^5^ and PolyPhen-2^9^), and three meta-predictors (REVEL^14^, GAVIN^15^ and ClinPred^16^) using a publicly available (‘open’) variant dataset and a clinically-relevant (‘clinical’) variant dataset. We show that the tools’ performance is heavily affected by the test dataset, and that all tools may perform worse than expected when classifying novel missense variants. By assessing the effect of a consensus-based approach, our results support the use of a single classifier when performing variant classification.

## 2. MATERIALS AND METHODS

### 2.1 Open Dataset

(n=8795, see **Figure S1A**) represents the typical training and validation dataset used during *in silico* predictor design and benchmarking. Positive (‘pathogenic’) variants were downloaded from ClinVar^12^ on 13^th^ November 2017 and subscription-based HGMD^28^ Professional release 2017.3; neutral (‘benign’) variants in OMIM^27^ morbid genes were downloaded from the gnomAD^11^ database (exomes only data v2.0.1). **ClinVar criteria:** Stringent criteria were used to increase the likelihood of selected variants being truly pathogenic. Missense SNVs with either ‘pathogenic’ and/or ‘likely pathogenic’ classification, multiple submitters and no conflicting submissions were included; variants with any assertions of ‘uncertain’, ‘likely benign’ or ‘benign’ were excluded. **HGMD Pro criteria:** Single nucleotide missense variants marked as disease-causing (‘DM’) were taken from HGMD Professional release 2017.3. **gnomAD criteria:** Missense SNVs with an overall minor allele frequency (MAF) between 1% and 5% were selected. These variants were deemed too common to be disease-causing but are not necessarily filtered out by next-generation sequencing pipelines depending on the MAF thresholds used. Chromosomal locations with more than one variant (multiallelic sites) were excluded. Any variants found to be present in the ‘pathogenic’ and ‘neutral’ datasets were removed from the both.

### 2.2 Clinical Dataset

(n=1766, see **Figure S1B** and **Supplemental Table S1**) more accurately reflects variants that might require classification in a clinical diagnostics laboratory following identification in an exome or genome sequencing pipeline. Variants were selected from three sources. **Group 1** (‘DDD’) consists of pathogenic (n=687) and benign (n=533) missense variants identified from 13,462 families in the Deciphering Developmental Disorders (DDD) study that have been through multiple rounds of variant filtering and clinical evaluation^26,29^. Variants were identified through exome sequencing and were reported to the patients’ referring clinicians for interpretation and confirmation in accredited UK diagnostic laboratories. All benign variants from this list were assessed as having no contribution towards the patient’s phenotype, and were present in either as heterozygotes in monoallelic genes or homozygotes in biallelic genes classified according to the Developmental Disorder Genotype-2-Phenotype database (DDG2P)^30^ (data accessed 17/10/2019). **Group 2** (‘Diagnostic’) consisted of pathogenic (n = 322) and benign (n=23) missense variants identified through Sanger sequencing, next-generation sequencing panel analysis or single gene testing in an accredited clinical diagnostic laboratory. Variants were manually classified according to the ACMG guidelines on variant interpretation^1^ on a 5-point scale (data accessed 23/04/2019). **Group 3** (‘Amish’) consisted of benign missense variants (n = 53) identified through a Community Genomics research study of 220 Amish individuals. Variants were identified through singleton exome sequencing and were classified as benign based on population frequencies and zygosity within this study. Two subgroups were manually selected and annotated based on inheritance pattern and disease penetrance; subgroup (i) consisted of variants in genes that cause a dominantly-inherited disorder with complete penetrance in childhood, for which the individual was clinically unaffected; this list was curated by a consultant in clinical genetics; subgroup (ii) consisted of variants in all other OMIM morbid genes (including those with incompletely penetrant dominant disorders and recessive and X-linked inheritance), with MAF>5% in the Amish cohort and MAF≤0.01% in gnomAD (data accessed 18/10/2019).

### 2.3 Transcript selection and variant annotation

For the open dataset, the canonical transcript was selected for each variant using the Variant Effect Predictor (VEP)^31^. For the clinical dataset, the HGMD Professional RefSeq transcript was used, unless absent from the database, in which case the MANE primary transcript was selected. Variants were annotated with variant cDNA and protein nomenclature in reference to the selected transcript. PolyPhen-2 and SIFT scores were annotated using VEP. REVEL and ClinPred scores were annotated using flat files containing precomputed scores for all possible single nucleotide substitutions, and in both cases, the combination of nucleotide position, nucleotide change and amino acid change was sufficiently unique to identify a single record, i.e. transcript selection did not affect the scores. GAVIN scores were generated through a batch submission to the GAVIN server.

### 2.4 Tool benchmarking

The performance of each of the tools was determined for both datasets. For SIFT, PolyPhen-2, REVEL and ClinPred, the output of the analysis was a numerical score between 0 and 1. Initially, all tools were analysed according to the criteria defined in their original publications, with the thresholds for pathogenicity being ≤0.05 for SIFT, ≥0.9 for PolyPhen-2 and ≥0.5 for ClinPred. For REVEL, where no threshold is recommended, a threshold of ≥0.5 was used. The categorical classification of GAVIN was used directly (“Benign”, “Pathogenic”; variants of uncertain significance (“VOUS”) were removed). A supplementary analysis was done for those tools with a numerical output (SIFT, PolyPhen-2, REVEL and ClinPred), to more accurately compare their performance. A unique threshold was selected for each tool to calculate the specificity when sensitivity was set to 0.9. In order to include GAVIN in this analysis, a third analysis was performed, whereby each tool’s specificity was measured when the threshold was adjusted to set the sensitivity identical to that of GAVIN.

## 3. RESULTS

### 3.1 Classification of variant sources

We compared the feature list of all tools benchmarked in this study (PolyPhen-2, SIFT, REVEL, GAVIN and ClinPred) and, in the case of the meta-predictors, the tools that they use as part of their algorithm (MPC^32^, MutPred^33^, VEST^34^, CADD^35^, DANN^36^, SNPEff^37^, FATHMM^38^, FitCons^39^ and MutationTaster^40^). Features were split into five broad categories: Conservation, Genetic variation, Functional evidence (nucleotide), Functional evidence (protein) and Amino acid properties (see **Figure 1** and **Supplemental Figure S2**). In general, the meta-predictors employ a wider variety of sources, and are less heavily reliant on conservation alone. CADD/DANN and FitCons, and by extension GAVIN and ClinPred, are the only predictors with features within the *Functional (nucleotide)* category and are therefore able to predict the pathogenicity of a variant in the context of its nucleotide change, regardless of whether there is a resultant amino acid change.

**Figure 1.**
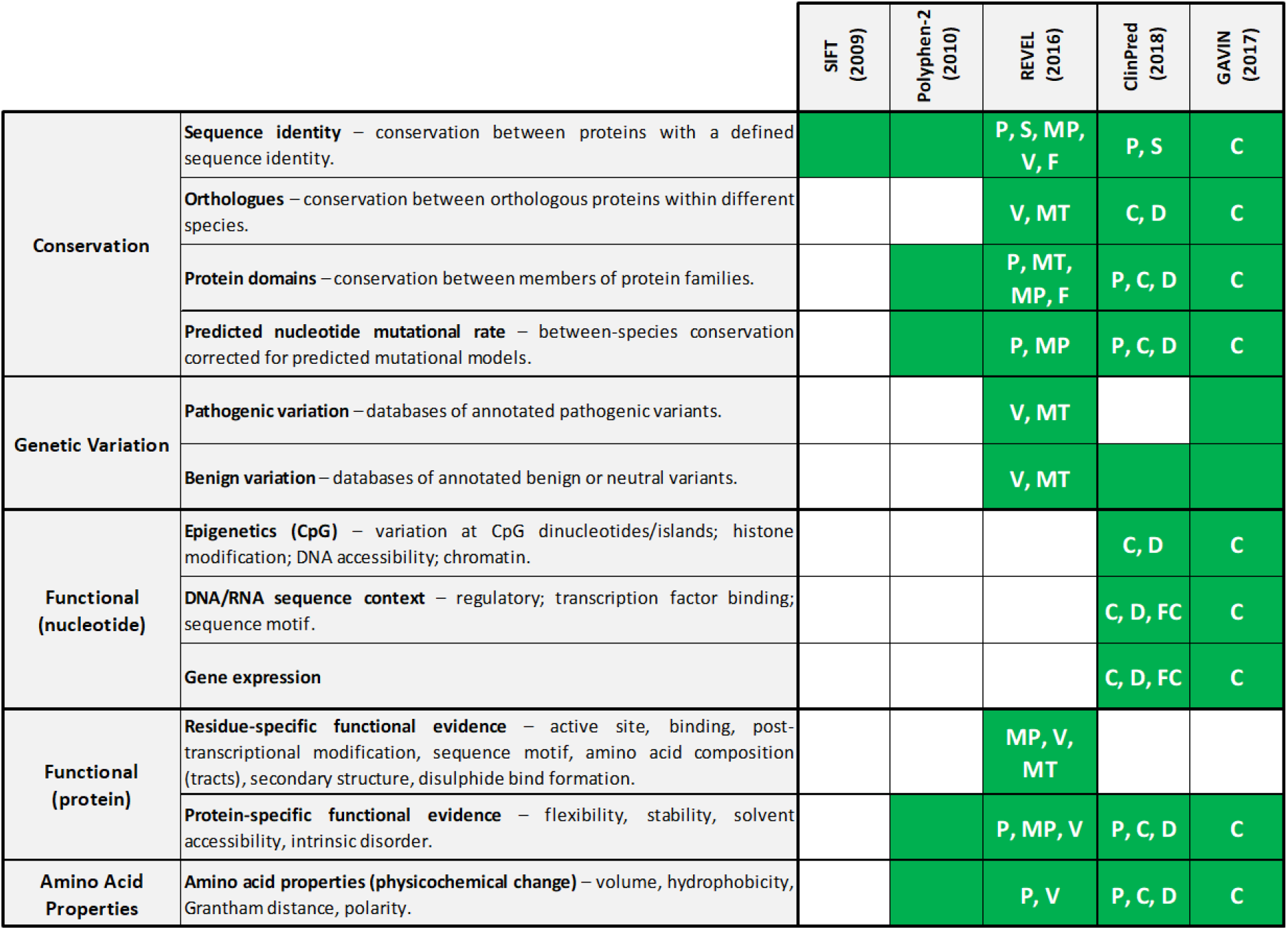
*In silico* pathogenicity predictor feature usage and source. Shading indicates that a category of evidence is utilised by the tool. Codes within each box indicate that the feature is inherited from another tool. Feature lists were taken from the tools’ original publications, supplementary materials and available online material. C – CADD; D – DANN; F – FATHMM; MP – MutPred; MT – MutationTaster; P – PolyPhen-2; S – SIFT; V – VEST; An extended version is shown in Supplemental Figure S2.

### 3.2 Benchmarking predictor performance for in the open and clinical datasets

Initially, each of the tools was benchmarked according to the threshold provided by the tools’ authors. This analysis involved a dichotomisation of scores with no intermediate range, see **Table 1**.

**Table 1.**
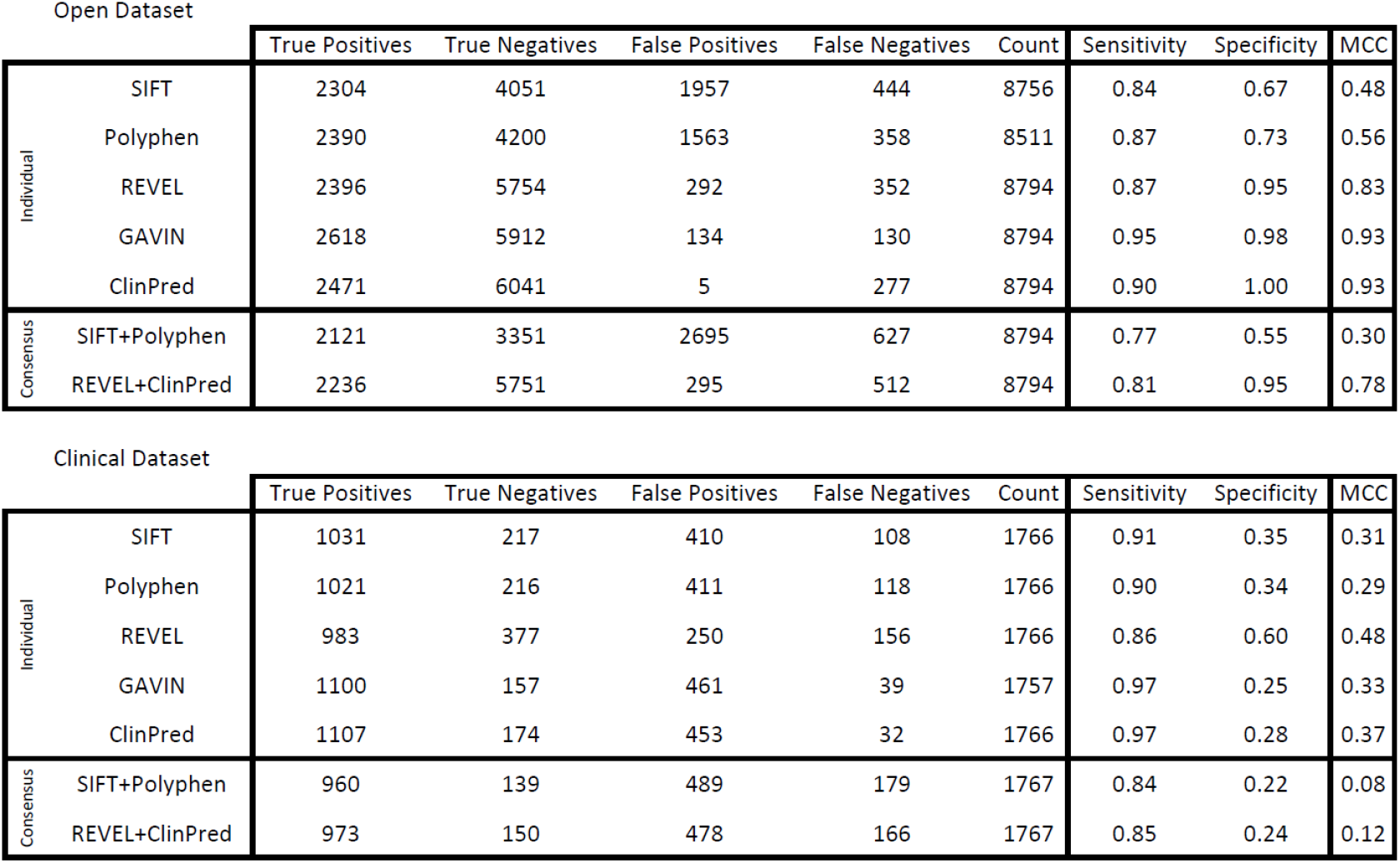
Results of variant classification for individual tool, and two consensus-based combinations, for datasets A, B and C. For consensus-based results non-concordant, where tools disagree on the classification, were considered incorrect.

| Open Dataset |  | True Positives | True Negatives | False Positives | False Negatives | Count | Sensitivity | Specificity | MCC |
| --- | --- | --- | --- | --- | --- | --- | --- | --- | --- |
| Individual | SIFT | 2304 | 4051 | 1957 | 444 | 8756 | 0.84 | 0.67 | 0.48 |
|  | Polyphen | 2390 | 4200 | 1563 | 358 | 8511 | 0.87 | 0.73 | 0.56 |
|  | REVEL | 2396 | 5754 | 292 | 352 | 8794 | 0.87 | 0.95 | 0.83 |
|  | GAVIN | 2618 | 5912 | 134 | 130 | 8794 | 0.95 | 0.98 | 0.93 |
|  | ClinPred | 2471 | 6041 | 5 | 277 | 8794 | 0.90 | 1.00 | 0.93 |
| Consensus | SIFT+Polyphen | 2121 | 3351 | 2695 | 627 | 8794 | 0.77 | 0.55 | 0.30 |
|  | REVEL+ClinPred | 2236 | 5751 | 295 | 512 | 8794 | 0.81 | 0.95 | 0.78 |

| Clinical Dataset |  | True Positives | True Negatives | False Positives | False Negatives | Count | Sensitivity | Specificity | MCC |
| --- | --- | --- | --- | --- | --- | --- | --- | --- | --- |
| Individual | SIFT | 1031 | 217 | 410 | 108 | 1766 | 0.91 | 0.35 | 0.31 |
|  | Polyphen | 1021 | 216 | 411 | 118 | 1766 | 0.90 | 0.34 | 0.29 |
|  | REVEL | 983 | 377 | 250 | 156 | 1766 | 0.86 | 0.60 | 0.48 |
|  | GAVIN | 1100 | 157 | 461 | 39 | 1757 | 0.97 | 0.25 | 0.33 |
|  | ClinPred | 1107 | 174 | 453 | 32 | 1766 | 0.97 | 0.28 | 0.37 |
| Consensus | SIFT+Polyphen | 960 | 139 | 489 | 179 | 1767 | 0.84 | 0.22 | 0.08 |
|  | REVEL+ClinPred | 973 | 150 | 478 | 166 | 1767 | 0.85 | 0.24 | 0.12 |

Matthews correlation coefficient (MCC) was calculated as follows:

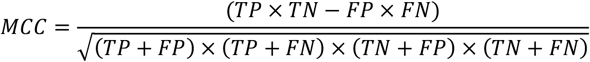

TP = True Positives; FP = False Positives; TN = True Negatives; FN = False Negatives;

The distribution of scores from SIFT, PolyPhen-2, REVEL and ClinPred is shown in **Figure 2** and ROC curves are shown in **Figure 3**. Of the tools with numerical outputs, ClinPred has the highest discriminatory power for the open dataset with an area under the ROC curve (AUC) of 0.993, while REVEL has the highest AUC for the clinical dataset (0.808). The two meta-predictors outperformed SIFT and PolyPhen-2 in both datasets. In agreement with tool author benchmarking^14–16^ the meta-predictors REVEL, ClinPred and GAVIN were highly proficient at classifying the variants in the open dataset, achieving sensitivities of 0.87, 0.90 and 0.95, and specificities of 0.95, 1.00 and 0.98, respectively. For variants in the clinical dataset, although the sensitivity each tool remained largely constant, the specificity of all tools dropped considerably. For REVEL, ClinPred and GAVIN, specificity is reduced to 0.62, 0.28 and 0.25, respectively **[Table 1]**.

**Figure 2.**
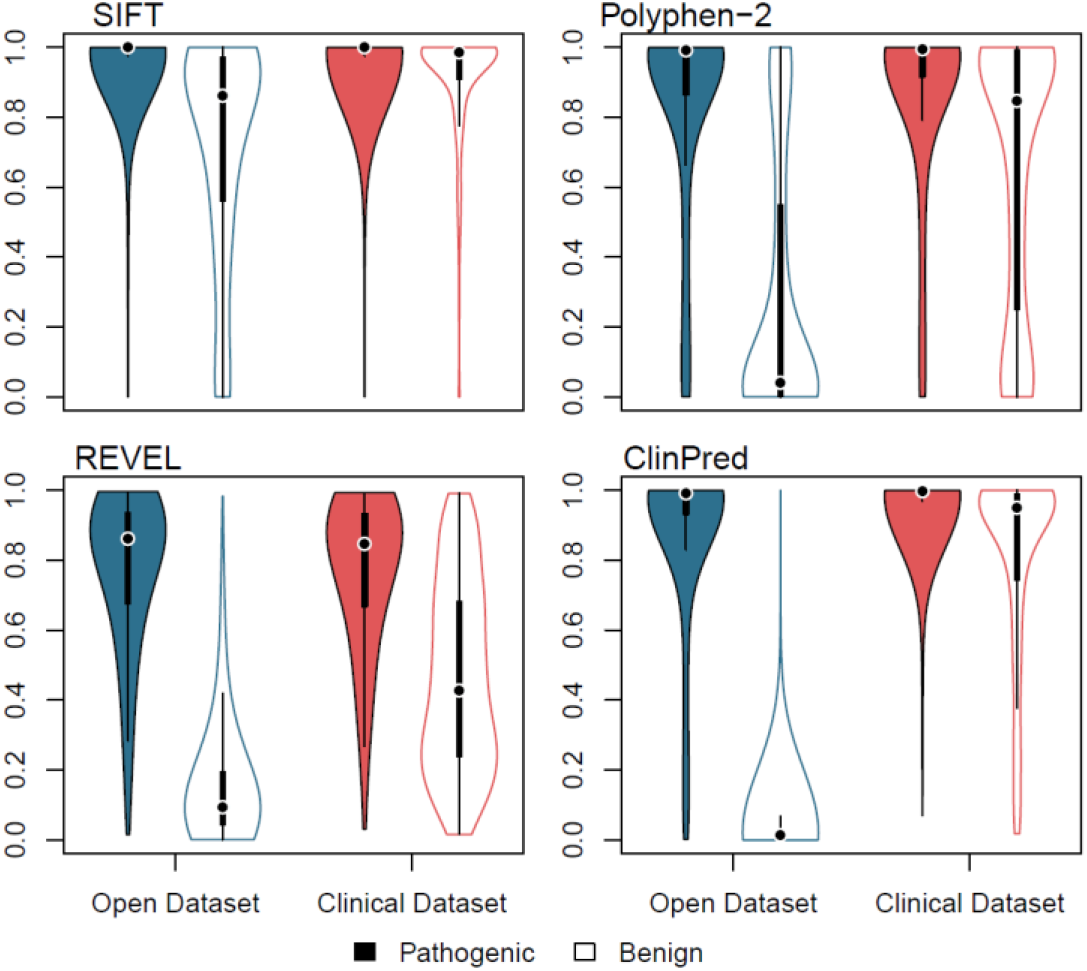
Violin plot showing variant scores for SIFT, PolyPhen-2, REVEL and ClinPred using two datasets. Open dataset – blue; clinical dataset – red; pathogenic variants – filled; benign variants – unfilled. Plot was generated in *R* using the ‘vioplot’ function in the ‘vioplot’ library. For ease of comparison, SIFT scores have been inverted.

**Figure 3.**
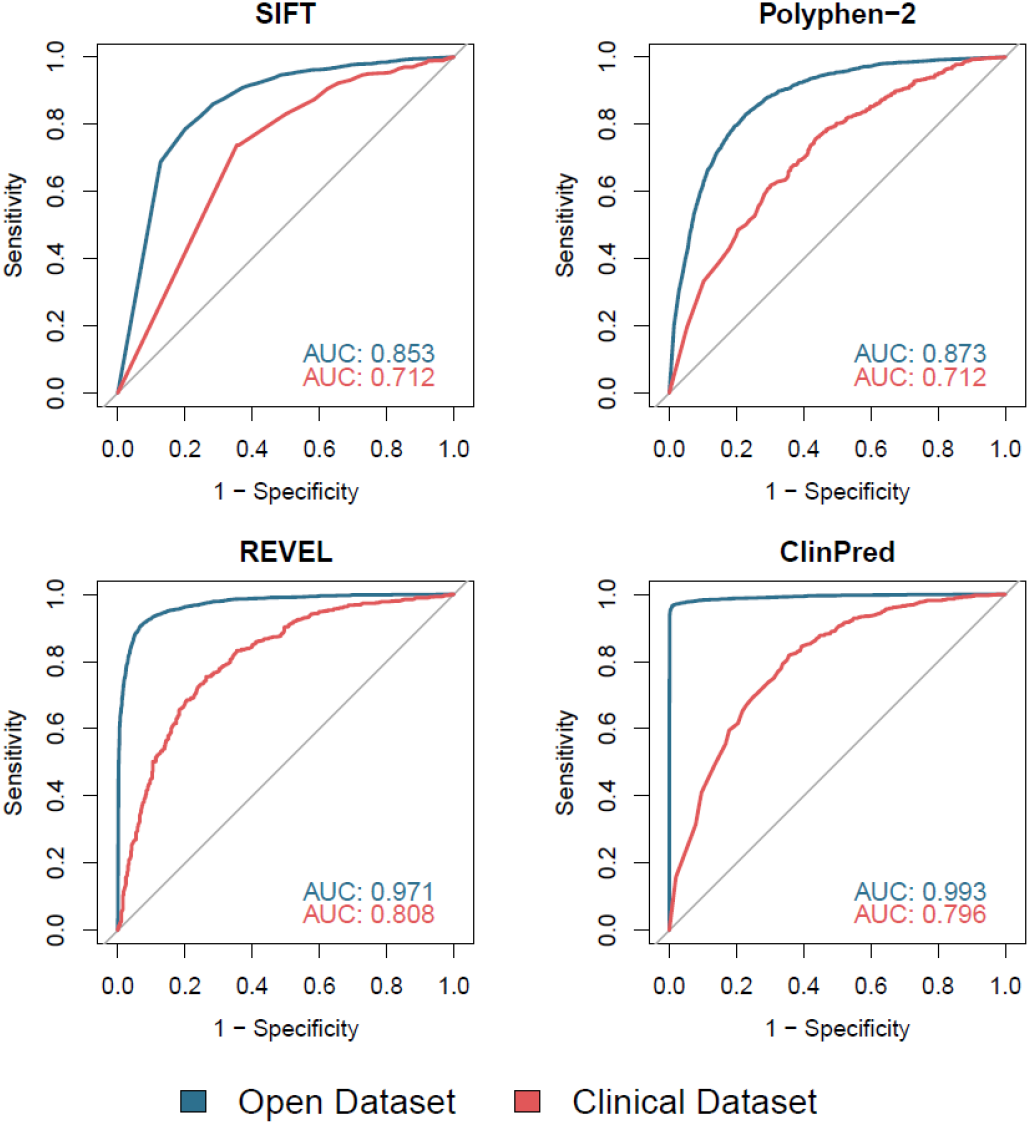
Receiver operating characteristic (ROC) curves for SIFT, PolyPhen-2, REVEL and ClinPred using two datasets. Open dataset – blue; clinical dataset – red. Generated in *R* using the ‘roc’ and ‘plot.roc’ functions in the ‘pROC’ library. Area under the ROC curve (AUC) was calculated in R using the ‘roc’ function. For ease of comparison, SIFT scores have been inverted.

It was apparent that the threshold suggested by the tools’ authors was not well-suited to both datasets, given the tools’ very high sensitivity but low specificity in the clinical dataset. In order to correct for this we performed a supplementary analysis for those predictors which gave a numerical output (SIFT, PolyPhen-2, REVEL and ClinPred). Here, a variable threshold was allowed for each tool to give a common sensitivity of 0.9 (i.e. pathogenic variation is called correctly 90% of the time). The threshold required to give a sensitivity of 0.9 in each tools is shown in **Table S2**. The specificity of each tool at the determined threshold is shown in **Figure S3**. When allowed a variable threshold the tools’ specificity increased significantly, with PolyPhen-2, SIFT, REVEL and ClinPred achieving a specificity of 0.67, 0.63, 0.93 and 0.99 for the open dataset, and 0.34, 0.33, 0.52 and 0.52 for the clinical dataset, respectively. In order to include GAVIN in this analysis, a third analysis was performed in which each tool was given a threshold to match the sensitivity achieved by GAVIN in each of the datasets. The specificity of all five tools is shown in **Figure S4**, and the sensitivity and threshold for each tool is shown in **Table S3**.

### 3.3 Use of individual tools versus a consensus-based approach between multiple tools

In accordance with current variant classification guidelines, we investigated the effect of performing a consensus-based analysis, using two commonly-used tools, SIFT and PolyPhen-2, and two meta-predictors, REVEL and ClinPred, to determine whether this combined approach has improved sensitivity/specificity over the individual tools. **Figure 4** shows the true concordance rate (variants classified correctly by both tools), false concordance rate (variants classified incorrectly by both tools) and discordance rate (variants for which the tools disagreed) for each of these tool pairings for the pathogenic and benign variants in both datasets. Within the clinically-relevant dataset, the tools are either falsely concordant or discordant for ~15% of pathogenic variants but ~77% of benign variants. The sensitivity and specificity of this approach is shown in **Table 1**. Use of a consensus-based approach introduces a third “discordance” category to the classification where no *in silico* evidence can be used, which applied to 24% and 16% of variants when considering the concordance of PolyPhen-2 and SIFT, and 8% and 23% when considering the concordance between REVEL and ClinPred, for the open and clinical datasets, respectively.

**Figure 4.**
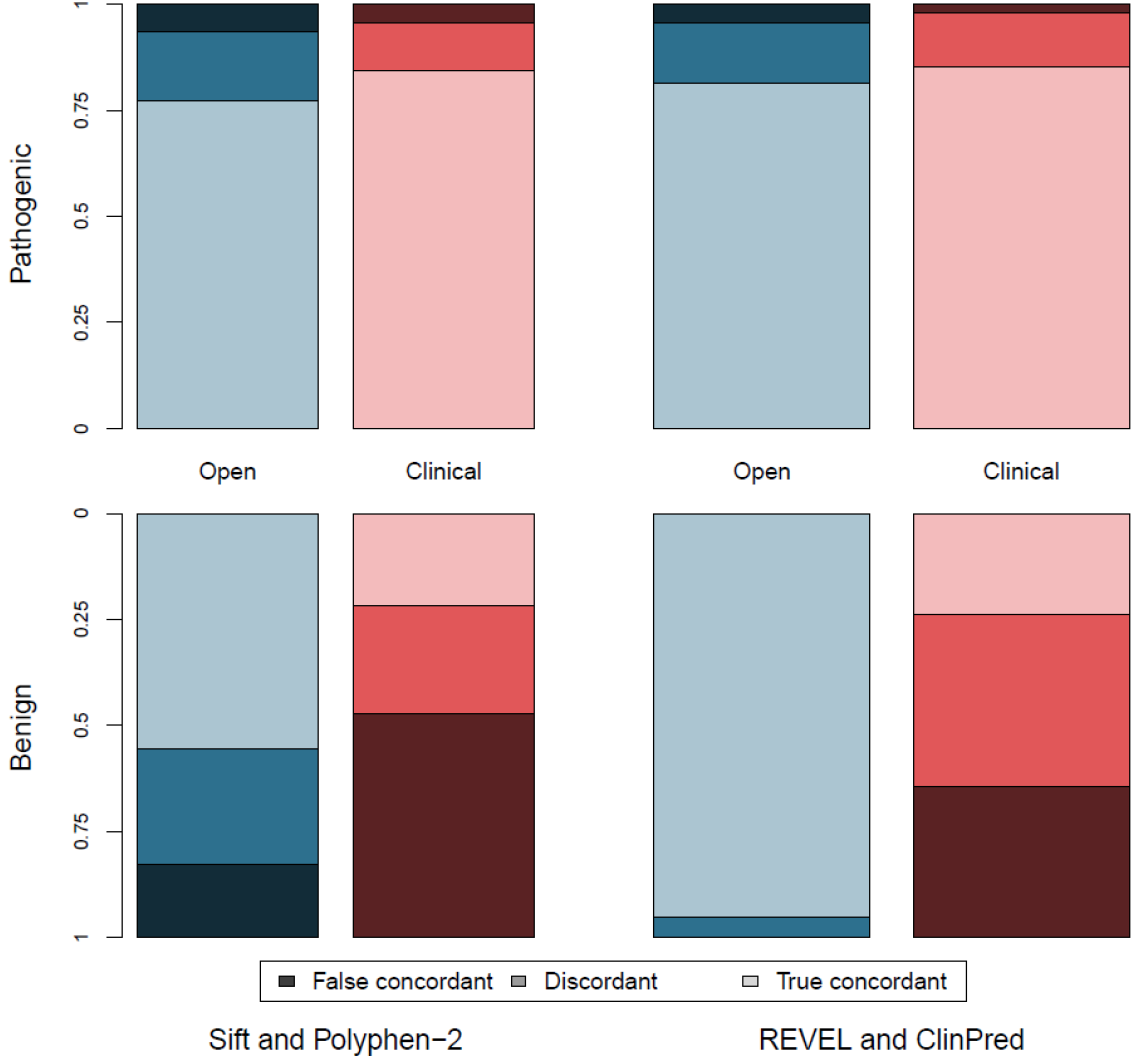
Concordance between tools separated by dataset and classification (pathogenic and benign). Open dataset – blue; clinical dataset – red; pathogenic variants – top graph; benign variants – bottom graph. True concordance indicates that the tools agree, and were correct. False concordance indicates that the tools agree but were incorrect. Discordance indicates that the tools disagreed on the classification.

## 4. DISCUSSION

We have compared the performance of five *in silico* pathogenicity predictors – two tools used routinely in variant classification (SIFT and PolyPhen-2) and three recently developed meta-predictors (REVEL, ClinPred and GAVIN) – using two variant datasets: an open dataset collated using the selection strategy commonly employed when benchmarking tool performance, and a clinically-representative dataset composed of rare and novel variants identified through high-throughput research and clinical sequencing and manual classification. Overall, the data herein show that meta-predictors have a greater sensitivity and specificity than the classic tools in both variant datasets. However, despite the increased accuracy of the meta-predictors, all tools performed substantially worse in the clinical dataset compared with the open dataset. This difference in tool performance illustrates the importance of considering the provenance of variants when benchmarking tools and how overfitting of a classifier to the training dataset can occur when increasingly large sets of variant features are utilised. Our analysis suggests that REVEL performs best when classifying rare variants routinely identified in clinical sequencing pipelines, with an AUC for our clinical dataset of 0.808, followed closely by ClinPred with an AUC of 0.796 **[Figure 3]** and with a higher specificity than GAVIN in a direct (albeit suboptimal) comparison **[Figure S4]**. While the REVEL team does not suggest a strict threshold for categorisation, in our analysis for the clinical dataset, a threshold of 0.43 gave a sensitivity of 0.9, and a specificity of 0.52, which is comparable to previous studies’ threshold of 0.5^16^.

Current guidelines on the classification of variants indicate that evidence should only apply when multiple tools are concordant^1^. However, the use of concordance introduces a third category to variants classification (discordance), where there is disagreement between tools and therefore the tools cannot be used as evidence to categorise the variant as either benign or pathogenic. Our data show that the use of concordance between multiple tools gives a lower sensitivity and specificity than the use of either of these tools in isolation, and furthermore that their performance is much below that of the meta-predictors.

As with all similar studies, we were limited by the availability of novel variants not present in online databases such as gnomAD. The use of under-represented and genetically isolated populations, such as the Amish, allowed for the identification of a number of novel benign variants and suggests that such populations may be a rich source for future studies. We also identified a number of both pathogenic and benign variants in a clinical population through a translational research study (DDD). While steps were taken to ensure that the benign variants attained from this group were indeed benign (all variant were present within either monoallelic genes, or in biallelic genes in a homozygous state, and were annotated by the referring clinician as having no contribution towards the patient’s clinical phenotype), nonetheless it cannot be guaranteed that the variants had no impact of protein function. The study highlights the need for improved data-sharing between clinical laboratories. While a number of online repositories exist for the sharing of rare pathogenic variants, no such resource is available for the sharing of rare benign variants.

The study supports the adoption of *in silico* meta-predictors for use in variant classification according to the ACMG guidelines, but recommends the use of a single meta-predictor over the application of a consensus-based approach. Each of the tools utilises different though heavily overlapping data sources and the feature list utilised by a tool should be carefully considered before the tool is utilised. Our results also suggests that tools that utilise gnomAD data directly may have low specificity when classifying rare or novel variants and that care should be taken when utilising these tools in conjunction with the ACMG guidelines. Although use of a meta-predictor tools offers notable advantages to the use of the previously available and widely adopted *in silico* tools, the remaining issues to be addressed before they can be used as more at a level greater than supporting evidence for clinical variant interpretation.

## Supplemental Materials

File S1: PDF file containing supplemental Figures S1, S2, S3, S5 and supplemental Tables S2 and S3.

File S2: Microsoft Excel file containing Supplemental Table S1.

## Web Resources

CADD: https://cadd.gs.washington.edu/

dbSNFP: https://sites.google.com/site/jpopgen/dbNSFP

GAVIN: https://molgenis20.gcc.rug.nl/menu/main/gavin-app

gnomAD: https://gnomad.broadinstitute.org/

HGMD Professional: https://portal.biobase-international.com/hgmd/pro/start.php

OMIM: https://www.omim.org/

PolyPhen-2: http://genetics.bwh.harvard.edu/pph2/

## Acknowledgements

We wish to thank all the patients and family members that participated in the study. We also thank Dr Michael Cornell and Dr Angela Davies, of the University of Manchester. We acknowledge funding from Wellcome [200990]. SE is a Wellcome Senior Investigator. The DDD study presents independent research commissioned by the Health Innovation Challenge Fund [grant number HICF-1009-003] a parallel funding partnership between the Wellcome Trust and the Department of Health, and the Wellcome Trust Sanger Institute [grant no. WT098051]. See Nature 2015;519:223-8 or www.ddduk.org/access.html for full acknowledgement.

