## Supplemental Materials for "Assessing performance of pathogenicity predictors using clinically-relevant variant datasets"

**Figure S1 - Flow diagram of selection and filtering steps used for the generation of the Open (A) and Clinical (B) datasets**

**Figure S2 - In silico pathogenicity predictor feature usage and source (extended).**

**Figure S3 - The specificity of SIFT, PolyPhen-2, REVEL and ClinPred for the Open (Blue) and Clinical (Red) datasets, with sensitivity set to 0.9.**

**Figure S4 - The specificity of SIFT, Polyphen-2, REVEL, ClinPred and GAVIN with sensitivity set to that of GAVIN**

**Table S2 - Threshold required to give an approximate sensitivity of 0.9.**

**Table S3 - Threshold required to give a sensitivity identical to that of GAVIN.**

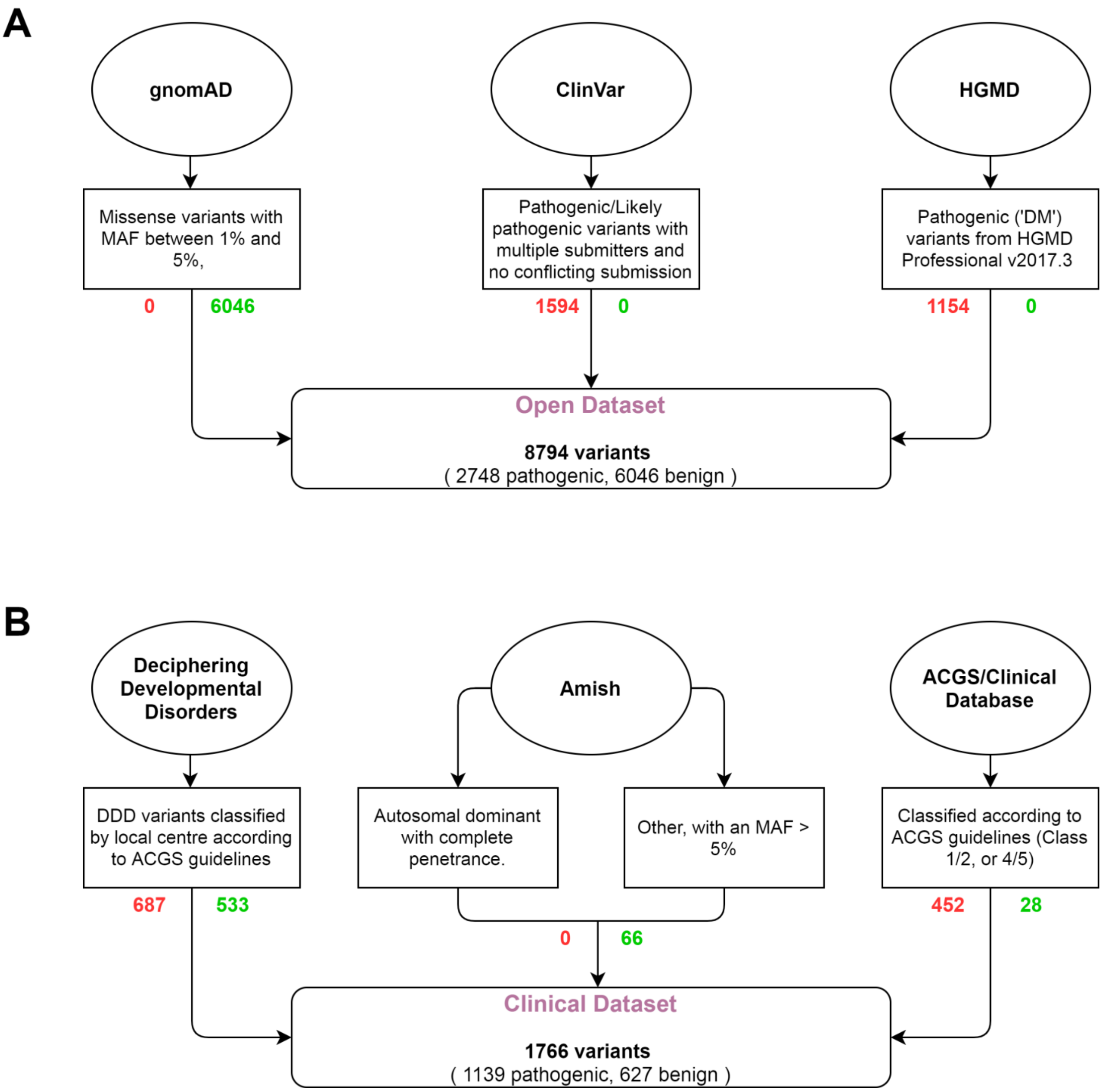


**Figure S1. Flow diagram of selection and filtering steps used for the generation of the Open (A) and Clinical (B) datasets.** Oval – variant source; Box – selection criteria; Rounded box – Dataset. Red text show number of pathogenic variants, green text shows benign variants.

| | | SIFT (2009) | PolyPhen-2 (2010) | REVEL (2016) | ClinPred (2018) | GAVIN (2017) | MutPred (2009) | MutationTaster (2010) | FATHMM (2013) | VEST (2013) | CADD (2014) / DANN (2015) |
|---|---|---|---|---|---|---|---|---|---|---|---|
| Conservation | **Sequence identity** – conservation between proteins with a defined sequence identity. | | | P, S, MP, V, F | P, S | C | | | | | P, S |
| | **Orthologues** – conservation between orthologous proteins within different species. | | | V, MT | C, D | C | | | | | |
| | **Protein domains** – conservation between members of protein families. | | | P, MT, MP, F | P, C, D | C | | | | | P |
| | **Predicted nucleotide mutational rate** – between-species conservation corrected for predicted mutational models. | | | P, MP | P, C, D | C | | | | | P |
| Genetic Variation | **Pathogenic variation** – databases of annotated pathogenic variants. | | | V, MT | | | | | | | |
| | **Benign variation** – databases of annotated benign or neutral variants. | | | V, MT | | | | | | | |
| Functional (nucleotide) | **Epigenetics (CpG)** – variation at CpG dinucleotides/islands; histone modification; DNA accessibility; chromatin. | | | | C, D | C | | | | | |
| | **DNA/RNA sequence context** – regulatory; transcription factor binding; sequence motif. | | | | C, D, FC | C | | | | | |
| | **Gene expression** | | | | C, D, FC | C | | | | | |
| Functional (protein) | **Residue-specific functional evidence** – active site, binding, post-transcriptional modification, sequence motif, amino acid composition (tracts), secondary structure, disulphide bind formation. | | | MP, V, MT | | | | | | | |
| | **Protein-specific functional evidence** – flexibility, stability, solvent accessibility, intrinsic disorder. | | | P, MP, V | P, C, D | C | | | | | P |
| Amino Acid Properties | **Amino acid properties (physicochemical change)** – volume, hydrophobicity, Grantham distance, polarity. | | | P, V | P, C, D | C | | P, C, D | | P | P |

**Figure S2. *In silico* pathogenicity predictor feature usage and source (extended).** Extended version of Figure 1 to include additional tools utilised by meta-predictors. Shading indicates that a category of evidence is utilised by the tool. Codes within each box indicate that the feature is inherited from another tool. Feature lists were taken from the tools' original publications, supplementary materials and available online material. P – PolyPhen-2; S – SIFT; MP – MutPred; V – VEST; C – CADD; D – DANN; MT – MutationTaster; F – FATHMM.

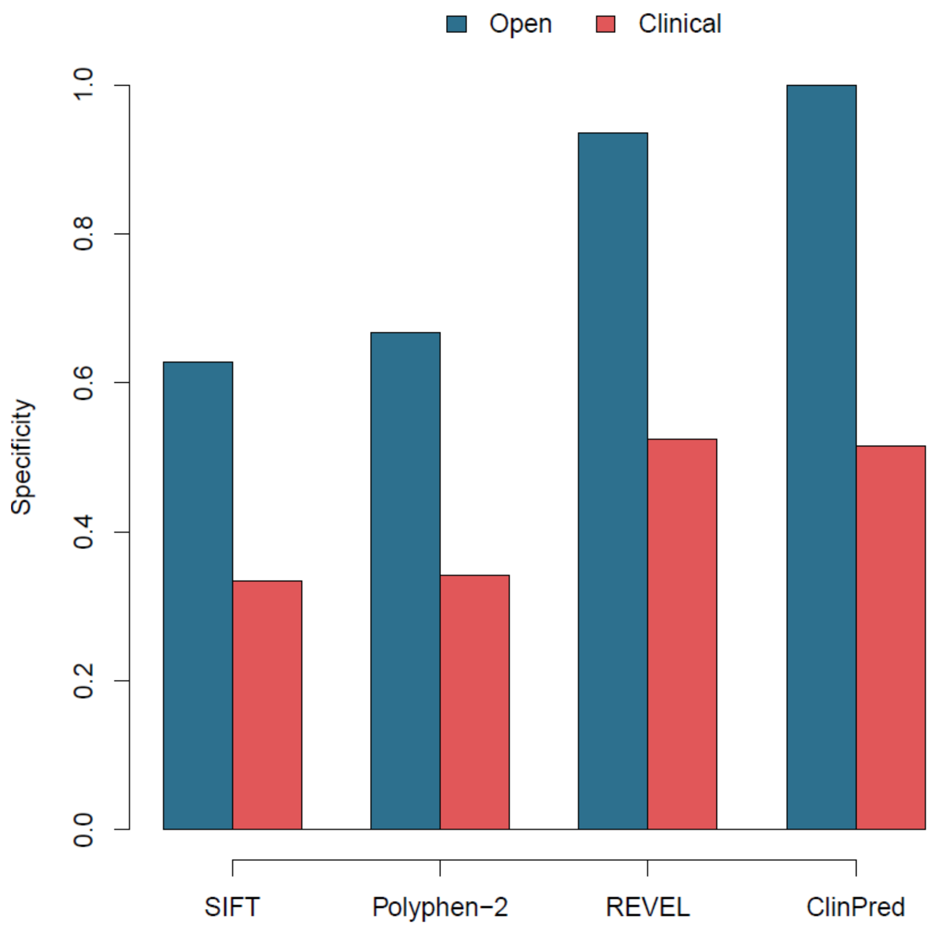


**Figure S3. The specificity of SIFT, PolyPhen-2, REVEL and ClinPred for the Open (Blue) and Clinical (Red) datasets, with sensitivity set to 0.9.** Thresholds for each tool required to give sensitivity of 0.9 are shown in Table S2.

| | Dataset | |
|---|---|---|
| | Open | Clinical |
| SIFT | ≤0.06 | ≤0.05 |
| Polyphen | ≥0.27 | ≥0.43 |
| REVEL | ≥0.44 | ≥0.43 |
| ClinPred | ≥0.50 | ≥0.91 |

**Table S2. Threshold required to give an sensitivity of 0.9.**

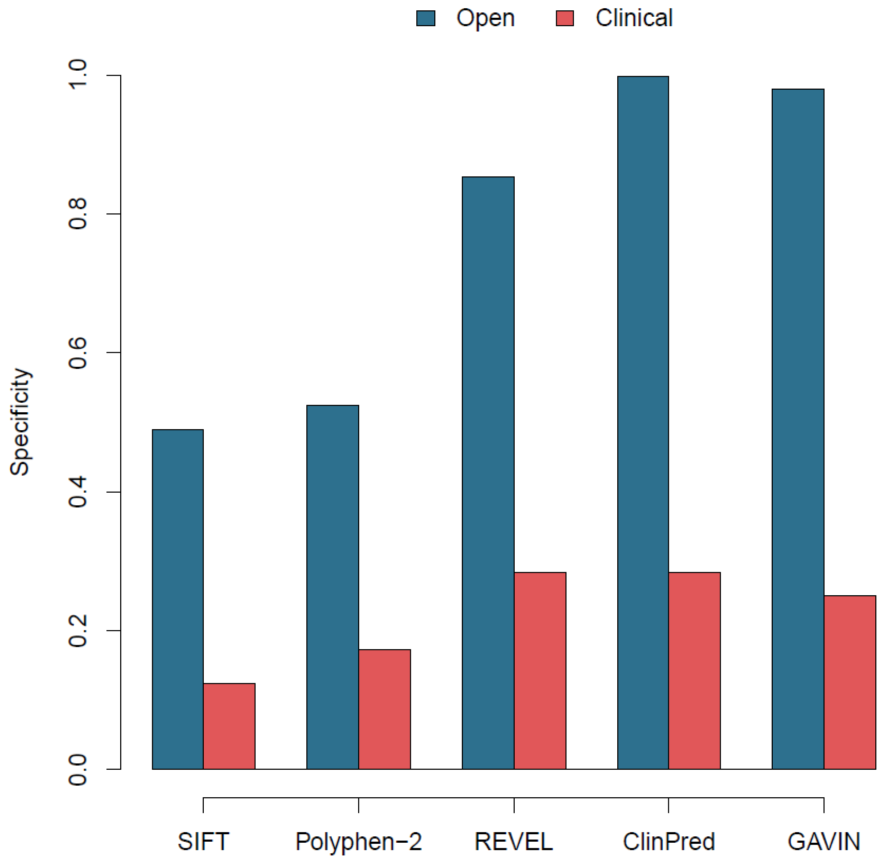


**Figure S4. The specificity of SIFT, Polyphen-2, REVEL, ClinPred and GAVIN for the Open (Blue) and Clinical (Red) datasets, with sensitivity set to that of GAVIN** (0.95 and 0.97 for the Open and Clinical datasets, respectively). Thresholds for each tool required are shown in Table S3.

| | Dataset | |
|---|---|---|
| | Open | Clinical |
| SIFT | ≤0.15 | ≤0.25 |
| PolyPhen-2 | ≥0.054 | ≥0.052 |
| REVEL | ≥0.29 | ≥0.24 |
| ClinPred | ≥0.17 | ≥0.52 |

**Table S3. Threshold required to give a sensitivity identical to that of GAVIN**
